# Invasion of the stigma by the pollen tube or an oomycete pathogen: striking similarities and differences

**DOI:** 10.1101/2023.07.19.549726

**Authors:** Lucie Riglet, Sophie Hok, Naïma Kebdani-Minet, Joëlle Le Berre, Mathieu Gourgues, Frédérique Rozier, Vincent Bayle, Lesli Bancel-Vallée, Valérie Allasia, Harald Keller, Martine Da Rocha, Thierry Gaude, Agnés Attard, Isabelle Fobis-Loisy

**Affiliations:** Laboratoire Reproduction et Développement des Plantes, Univ Lyon, ENS de Lyon, UCB Lyon1, CNRS, INRA, F-69342 Lyon, France; Sainsbury Laboratory, University of Cambridge, Cambridge, United Kingdom; INRAE, CNRS, Université Côte d’Azur, UMR1355-7254, ISA, 06903 Sophia Antipolis, France; Bayer Crop Science France, 14, impasse Pierre Baizet CS 99163, F-69263 Lyon, France; Unité de Bordeaux, Bordeaux Imaging Center, 146 rue Lèo Saignat CS 61292, F-33076 Bordeaux; Carl Zeiss SAS, 15 avenue Edouard Belin, F-92500 Rueil-Malmaison, France

## Abstract

The epidermis is the first barrier that protects organisms from surrounding stresses. Similar to the hyphae of filamentous pathogens that penetrate and invade the outer tissues of the host, the pollen germinates and grows a tube within epidermal cells of the stigma. Early responses of the epidermal layer are therefore decisive for the outcome of these two-cell interaction processes. Here, we aim at characterizing and comparing how the papillae of the stigma respond to intrusion attempts, either by hypha of the hemibiotrophic oomycete root pathogen, *Phytophthora parasitica* or by the pollen tube. We found that *P. parasitica* spores attach to the papillae and hyphae subsequently invade the entire pistil. Using transmission electron microscopy, we examined in detail the invasive growth characteristics of *P. parasitica* and found that the hypha passed through the stigmatic cell wall to grow in contact with the plasma membrane, contrary to the pollen tube that advanced engulfed within the two cell wall layers of the papilla. Further quantitative image analysis revealed that the pathogen and the pollen tube trigger reorganization of the endomembrane system (trans Golgi network, late endosome) and the actin cytoskeleton. Some of these remodeling processes are common to both invaders, while others appear to be more specific showing that the stigmatic cells trigger an appropriate response to the invading structure and somehow can recognize the invader that attempts to penetrate.

## Introduction

The epidermis is the outermost cell layer of plants that is in direct contact with the environment. epidermal cells have to promptly react to mediate the most relevant responses. Invaders can be infection structures such as hyphae of fungi or oomycetes but also reproductive structures like pollen tubes. The first contact between infection hyphae and epidermal cells is decisive for the outcome of the interaction: disease or resistance. Similarly, the first interaction that occurs between invading pollen tubes and the epidermal cells of the stigma (papillae) is crucial for successful reproduction. In both cases, a fine-tuned dialog is established at early stages of the interaction between the host and the invader and is critical for the result of these two cell-cell interaction systems.

Many points of convergence between pathogen defense and pollen recognition have already led some authors to suggest that the two processes share common origins (Nasrallah, 2005; Kodera et al., 2021). Briefly, (i) the fungal/oomycete hyphae and the pollen tube are tip growing cells that secrete cell wall-degrading enzymes to weaken and penetrate the host surface layer (Chapman and Goring, 2010; Kebdani et al., 2010; Blackman et al., 2014), (ii) subcellular reorganization of organelles and cytoskeleton occurs in epidermal cells at the penetration sites (Takemoto et al., 2003; Hardham, 2007; Iwano et al., 2007; Samuel et al., 2009; Samuel et al., 2011), (iii) both hyphae and pollen tubes take up resources from invaded cells for their growth. Moreover, plant receptor-like kinases are involved in the two processes, and sometimes can be common. The best example is the Feronia receptor, which acts as a scaffold for the assembly of the immune-receptor complex and regulates the pathogen-elicited burst of Reactive Oxygen Species (ROS) (Stegmann et al., 2017). It also controls the changes of ROS status in stigmatic cells allowing the pollen to germinate a tube (Liu et al., 2021). A transcriptomic analysis also predicted components of the pattern-triggered immunity to be activated in the stigma upon pollination (Kodera et al., 2021). Similarly, a study in which the transcriptome of pollinated pistils was compared to *Fusarium graminearum* infected ones revealed that similar groups of genes were overexpressed in pistil responding to pollen tubes or hyphae intrusions (Mondragón-Palomino et al., 2017). This study was conducted at late stage of interaction and is the only one, involving a common host tissue, the pistil, to compare reproductive and immune responses. So far, a detailed comparison of the cellular responses to intrusion and early growth of these two types of invasive organisms has never been carried out. Here, we identified the *A. thaliana* stigma as a common “host” for supporting pollen tube growth and infection with an oomycete pathogen. We show that *Phytophthora parasitica*, a hemibiotrophic telluric pathogen, attaches to and penetrates the papillae of the stigmatic epidermis before colonizing the entire pistil. Using transmission electron microscopy (TEM), we examined the invasive growth features of *P. parasitica* within the papillae and compared them with those of the pollen tube. Using Arabidopsis lines expressing fluorescent tagged-proteins for subcellular localisation studies, we found that both pathogen and pollen tube trigger cytoskeletal and endomembrane reorganization following intrusion. Taken together, our results show that stigma cells respond to *P. parasitica* invasion in a manner similar to plant cells that are natural targets of oomycete infection, and that some features are different from the stigma cell response to pollen tube invasion.

## Results

### *P. parasitica*, but not *H. arabidopsidis* colonizes the pistil

To investigate whether oomycete pathogens are able to break the stigmatic barrier and infect *Arabidopsis thaliana* pistils, we selected two oomycete species from different genera with different life styles and host ranges. *P. parasitica,* is a hemibiotrophic root pathogen with a wide host range including *A. thaliana* (Attard et al., 2010) and *Hyaloperonospora arabidopsidis*, is an obligate biotrophic foliar pathogen with *A. thaliana* as its sole host. On their natural host organs, both oomycetes penetrate within the first four hours after infection (hai). Mobile zoospores and immobile conidiospores from *P. parasitica* (Figure 1A) and *Hpa* (Figure 1B), respectively, emit a germ tube on the plant surface that forms a swelling structure (appressorium) dedicated to breach the epidermis through a penetrating hypha (Attard et al., 2010; Kebdani et al., 2010; Boevink et al., 2020).

**Figure 1.**
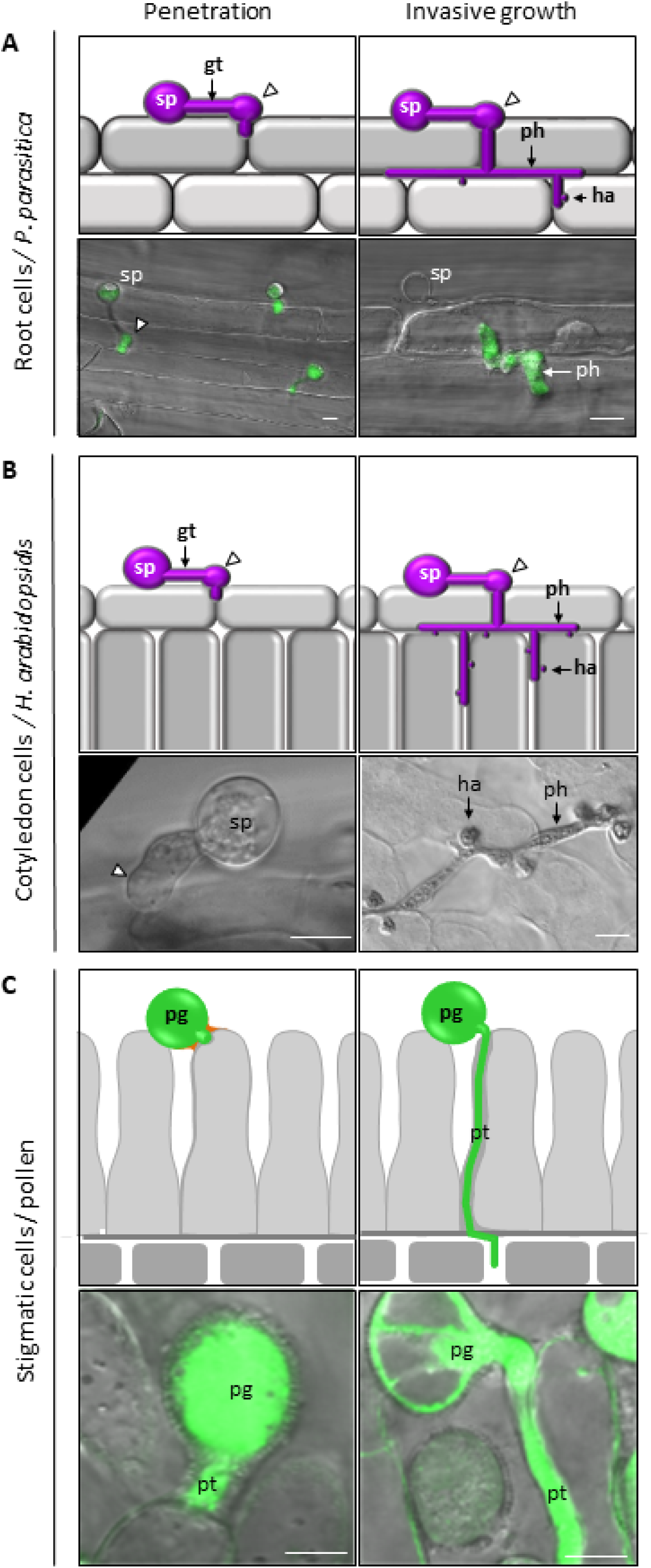
Invasion features of *P. parasitica*, *H. arabidopsidis*, and *A. thaliana* pollen tube in root, leaf, and stigma, respectively. A, Arabidopsis root infected with the oomycete *P. parasitica*. Schematic representations (upper row) and merged CLSM images between the green channel and bright-field of *P. parasitica* strain pCL380-GFP:GUS expressing a cytoplasmic GFP marker (lower row). The zoospore (sp) germinates a germ tube (gt) that grows along the plant surface and forms an appressorium (white arrow head) to penetrate the epidermis. Penetration starts 30 min after infection (left panel). The penetrating hypha (ph) grows intercellularly and develops haustoria (ha) (6 hai, right panel). B, Arabidopsis leaf infected with the oomycete *H. arabidopsidis*. Schematic representations (upper row) and bright-field CLSM images (lower row). The conidiospore (sp) germinates a germ tube (gt) that grows along the plant surface and forms an appressorium (white arrow head) to penetrate the epidermis (8 hai, left panel). The penetrating hypha (ph) grows intercellularly and develops haustoria (ha) (18 hai, right panel). C, Arabidopsis stigma with germinating pollen. Schematic representations (upper row) and merged CLSM images between green fluorescence and bright-field of a pollen grain and a pollen tube expressing the GFP marker (pLAT52-GFP Line; lower row). Rapidly after pollen capture, a specific structure, called the foot (depicted in orange), is formed at the pollen-papilla interface. A pollen tube (pt) emerges from the grain, passes through the foot and penetrates the papilla CW (dark grey layer) 12 map (left panel). The pollen tube grows inside the papilla CW towards the basis of the stigma (right panel). Bars represent 10 µm.

We applied conidiospores of *H. arabidopsidis* to the stigma by gently rubbing Arabidopsis leaves with sporulating conidiophores over the pistil surface from late stage 12 floral buds (before anthesis, Smyth et al., 1990). Four hours after inoculation, spores started to germinate a germ tube that grew around papillae (Figure 2A) but no appressorium were observed, suggesting that *H. arabidopsidis* does not manage to penetrate the stigma epidermis. *P. parasitica* produces zoospores that are motile and swim towards roots under natural conditions. To infect pistil tissues, we dipped either entire flower buds or naked pistils into a suspension of motile zoospores from a *P. parasitica* strain, which conditionally expresses a Green Fluorescent Protein and ß-glucuronidase (GFP:GUS) fusion protein upon zoospore germination with an expression level highly increasing during penetration of plant tissue (Attard et al., 2014). In both cases, zoospores preferentially accumulated at the stigma surface and penetrate the papillae, as revealed by the high expression of the GUS reporter (Figure 2, B and C). A preference for specific host tissues is also observed on roots, where zoospores expressing the GUS reporter aggregate around the elongation zone (Figure 2D). Twenty-four hours after infection, we observed GFP-labeled *P. parasitica* hyphae penetrating the pistil (Figure 2E). In contrast to pollen tubes, whose elongation was restricted to the central transmitting tract (Figure 2F), *P. parasitica* hyphae invaded the entire pistil body. Our observations show that the root pathogen *P. parasitica* (contrary to the leaf pathogen *H. arabidopsidis*) is able to overcome the stigmatic barrier and invade the pistil, although the stigmatic epidermis is not its natural host tissue.

**Figure 2.**
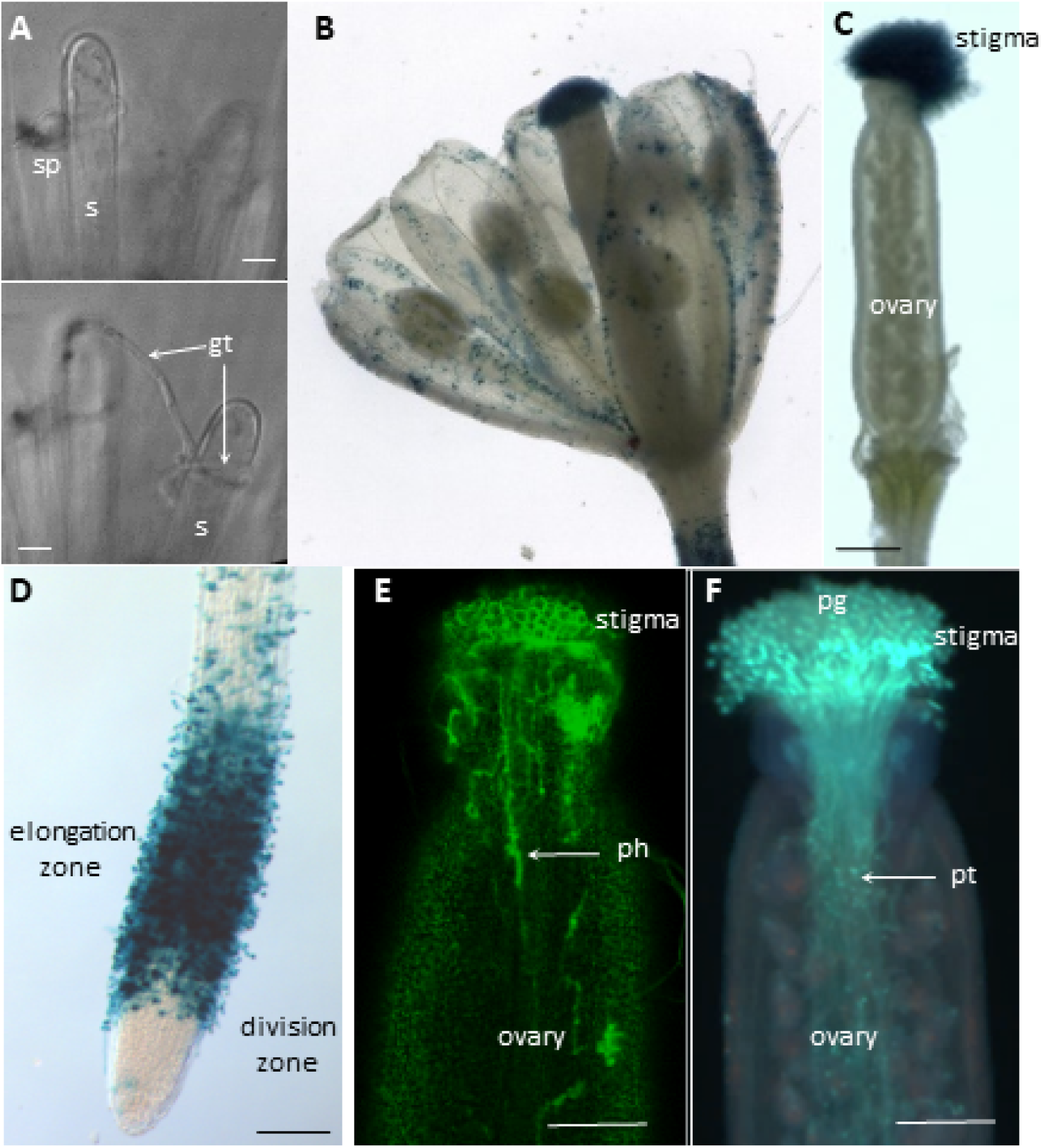
*P. parasitica* but not *H. arabidopsidis* invades *A. thaliana* pistil. A, *H. arabidopsidis* spores (sp) were deposited on the stigma surface of mature pistils and observed by CLSM. The upper and lower images are both extracted from the same Z-stack showing a spore and a germ tube (gt) growing around the papilla cells (s) without penetration, 18 hai. Bars represent 10 µm. B and C, The entire flower bud (B) or the naked pistil (C) were dipped in a suspension of GUS-expressing zoospores of *P. parasitica* (strain *pCL380-GFP:GUS*) and observed by transmission light microscopy. Zoospores preferentially attached to the stigma epidermis, three hai. D, *A. thaliana* roots dipped in a suspension of *P. parasitica* strain *pCL380-GFP:GUS* and observed by TEM. Zoospores preferentially attached to the elongation zone of the root, three hai. E, Entire flower buds were dipped in a suspension of GFP-expressing zoospores of *P. parasitica* and observed by CLSM. Median longitudinal optical section of the entire pistil. Growing hyphae were detected inside the pistil tissues 24 hai. F, Pollinated pistils stained with aniline blue and observed under epifluorescence microscopy. Six hours after pollen grain (pg) deposition at the stigma surface, pollen tubes (pt) were present within the central transmitting track. Bars in B-E represent 100 µm. Each experiment was repeated at least three times.

### *P. parasitica* forms appressoria for penetration and induces a PM-derived membrane around the invading hyphae

To further characterize *P. parasitica* infection, we compared the early stages of infection occurring at the epidermis of stigma and roots. One hour after pistil inoculation, zoospores developed a germ tube that grew on the papilla surface (Figure 3A). At the extremity of this germ tube a swelling appressorium-like structure is formed (white arrow head) and penetrating hyphae grew into the papilla cells (Figure 3B). We used an Arabidopis line expressing the GFP-tagged plasma membrane (PM) marker LTI6b in stigma (Rozier et al., 2020) to monitor papilla PM remodeling during infection. Four hai, a LTI6b-labeled membrane enclosed the invading *P. parasitica* hypha (Figure 3, C-E). Because little is known about the behavior of the PM in epidermal root cells invaded by *P. parasitica*, we examined the fate of the PM in root cells that undergo hyphal penetration using an Arabidopsis line expressing the RFP-tagged PM aquaporin AtPIP2A in root. Similar to what has been observed on the stigma zoospores at the root epidermis emitted a germ tube, formed an appressorium to penetrate the host epidermis and entered the cells (Figure 3, F-H). Inside the cells, a structure labeled with the PM marker encased the penetrating hyphae (Figure 3, I-K). Taken together, our observations suggest that *P. parasitica* uses comparable infection processes to invade papillae and root cells.

**Figure 3.**
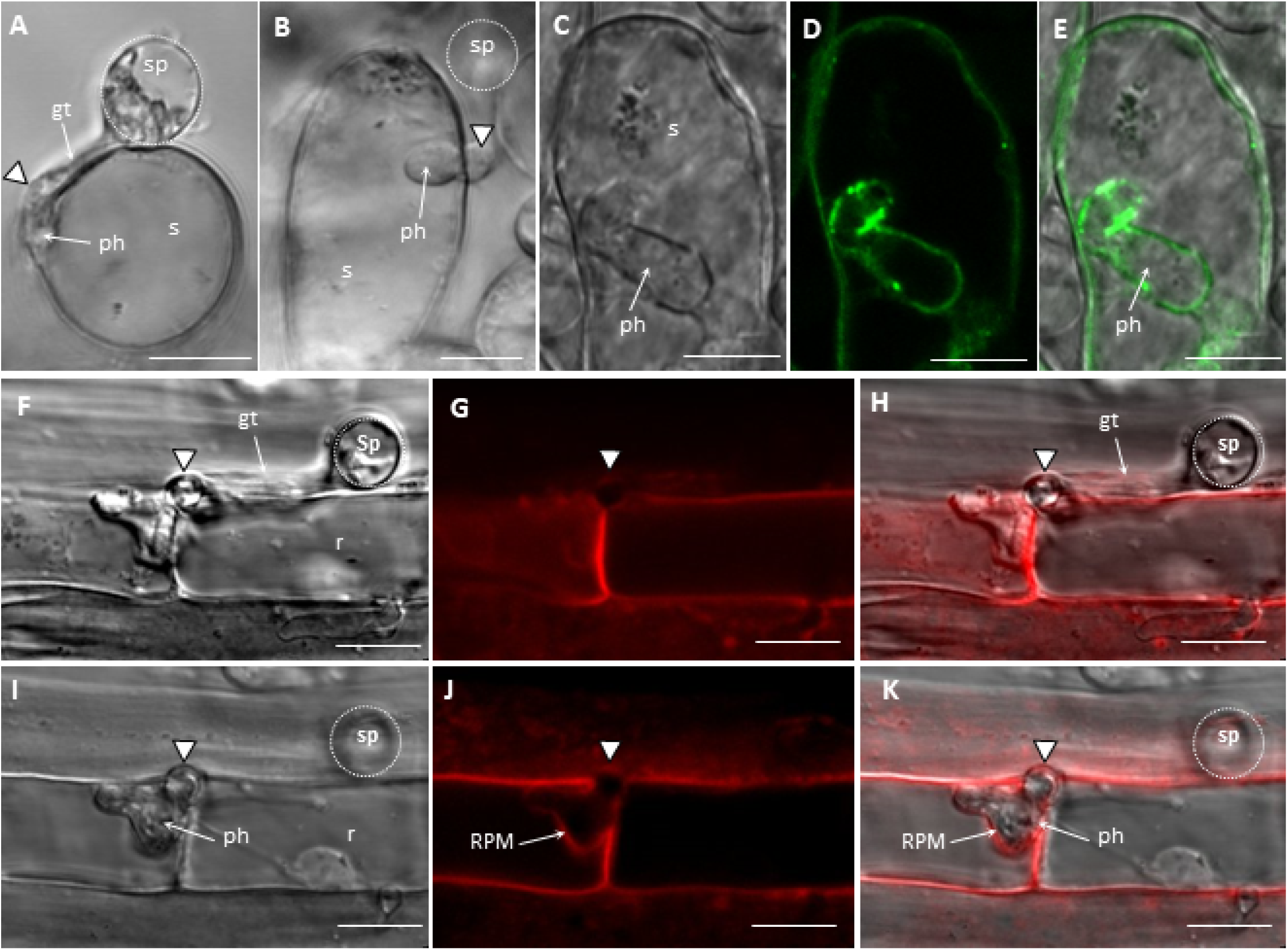
Appressorium-mediated penetration of the pistil and the root epidermis by *P. parasitica*. A, Transversal optical sections of a zoospore (sp, delimited by a white dashed line) of *P. parasitica* germinating a germ tube (gt) on the stigmatic cell surface, one hai. The germ tube forms an appressorium-like structure at its extremity (white arrow head). B, Longitudinal optical sections of *P. parasitica* infecting a papilla cell. The penetrating hypha (ph) emerges from an appressorium (white arrow head). C, D and E. An inoculated papilla expressing a GFP-tagged PM marker (LTI6b) four hai. A Lti6b-labeled membrane encircles the penetrating hypha, as observed in bright field (C), green fluorescence (D) and the merged image (E). F to K, Arabidopsis root expressing a RFP-tagged PM aquaporin *At*PIP2A upon invasion by *P. parasitica*. The roots were dipped in a zoospore suspension for one hour and the epidermis was analyzed by CLSM. The images show optical sections of the same infection site, with F, G and H focused on the root (r) surface, and I, J and K on the cell interior. The zoospore (sp) germinates a germ tube that differentiates an appressorium (white arrow head) to penetrate between two adjacent cells. Inside the epidermis, the *At*PIP2A-labeled membrane surrounds the penetrating hypha. Left column, bright field, middle column, RFP channel, and right column, merged channel. Bars represent 10 µm. Each experiment was repeated at least three times.

### Different mechanisms for penetration of stigmatic cells by pollen tube or infectious hyphae

To better analyze the invader/host cell interface, we performed TEM analyses. Three hours after inoculation with *P. parasitica,* the stigma cuticle and the stigma cell wall (CW) were not visible anymore beneath the appressorium-like structure at the papilla surface (Figure 4, A and C), strongly suggesting that both layers have been digested at the penetration site. Inside the papilla, the hypha was found between the cuticle and the CW with the stigmatic CW partially digested (Figure 4, A-D) or totally embedded in the stigmatic cytoplasm (Figure 4E). Because cell penetration is not synchronous, we assume that gradual CW digestion from partial (Figure 4C) to complete (Figure 4E) might correspond to different stages of infection.

**Figure 4.**
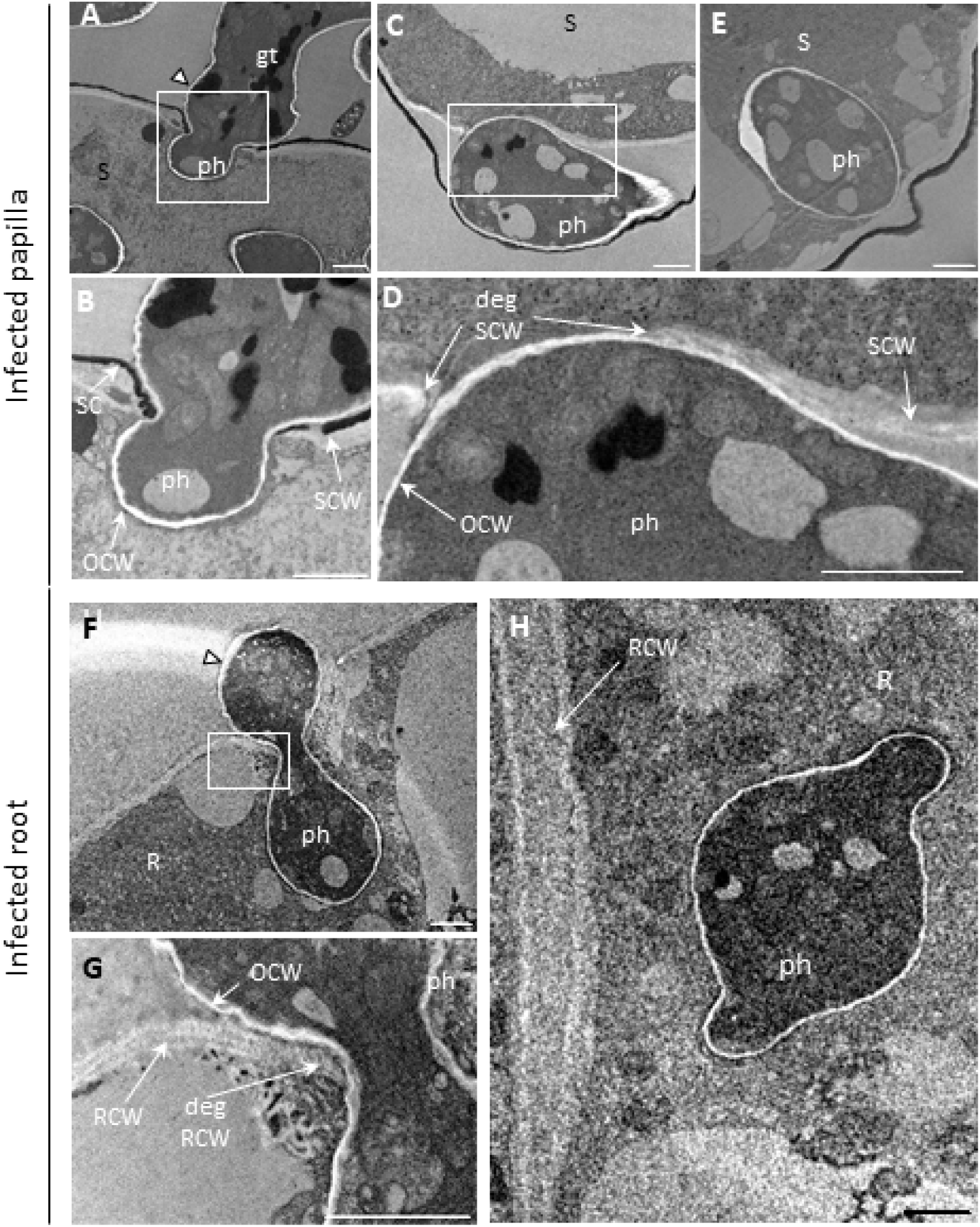
Invasion by *P. parasitica* hypha depends on the host epidermis. A to E, Stigmatic cell infected with *P. parasitica* observed by TEM, three hai. A, The germ tube (gt) emitted from the zoospore is located outside the papilla (s). The extremity of the germ tube forms an appressorium-like structure (white arrow head). A penetrating hypha (ph) enters the host cell. B, Magnification detail depicted by the white square in A showing the stigmatic cuticle (SC; electron dense black layer) and the stigmatic CW (SCW) digested at the penetration site. The oomycete CW (OCW) appears as an electron transparent white layer. C, The hypha locates between the stigmatic cuticle and the stigma CW (SCW). D, Magnification detail depicted by the white square in C showing the stigmatic CW degradation (degSCW) occurring at the contact area with the hypha. E, *P. parasitica* penetration hypha (ph) embedded in the stigmatic cytoplasm (s). F to H, Root cells (R) infected with *P. parasitica,* as observed by TEM, three hai. F, An appressorium (white arrowhead) is visible at the root surface and a penetration hypha (ph) inside the host cell. G, Detailed view depicted by the white square in F showing degradation of the root CW(deg RCW). H, A penetration hypha embedded in the root cytoplasm. Bars represent 1 µm. Each experiment was repeated at least three times.

We then compared the infection process of stigma with natural root infection. Three hai, the root CW was digested beneath the appressorium at penetration sites (Figure 4, F and G). As previously observed for infected papillae, we found *P. parasitica* hyphae embedded in the root cell cytoplasm (Figure 4H).

Similarly, we compared papilla infection with pollination. Rapidly after a compatible pollen grain comes in contact with a stigmatic cell, proteins and lipids from both cell surfaces fused to form a hydrophilic environment (called the foot) essential for pollen acceptance (Figure 1C; Chapman and Goring, 2010). This contact is followed by pollen hydration and emission of a pollen tube (germination) that invades the stigmatic cell to grow towards the stigma basis (Figure 1C). When analyzing papilla cells by TEM, we found that the pollen tube breached the cuticle layer 30 minutes after pollination (map, Figure 5, A and B). In contrast to hyphal penetration, we did not observe a complete digestion of the stigmatic CW by the pollen tube. Rather, the pollen tube grew between the inner and outer CW layers of the papillae (Figure 1C, Figure 5, C-E). This form of invasive pollen tube growth appears to be characteristic for *Brassicaceae* species (Riglet et al., 2020).

**Figure 5.**
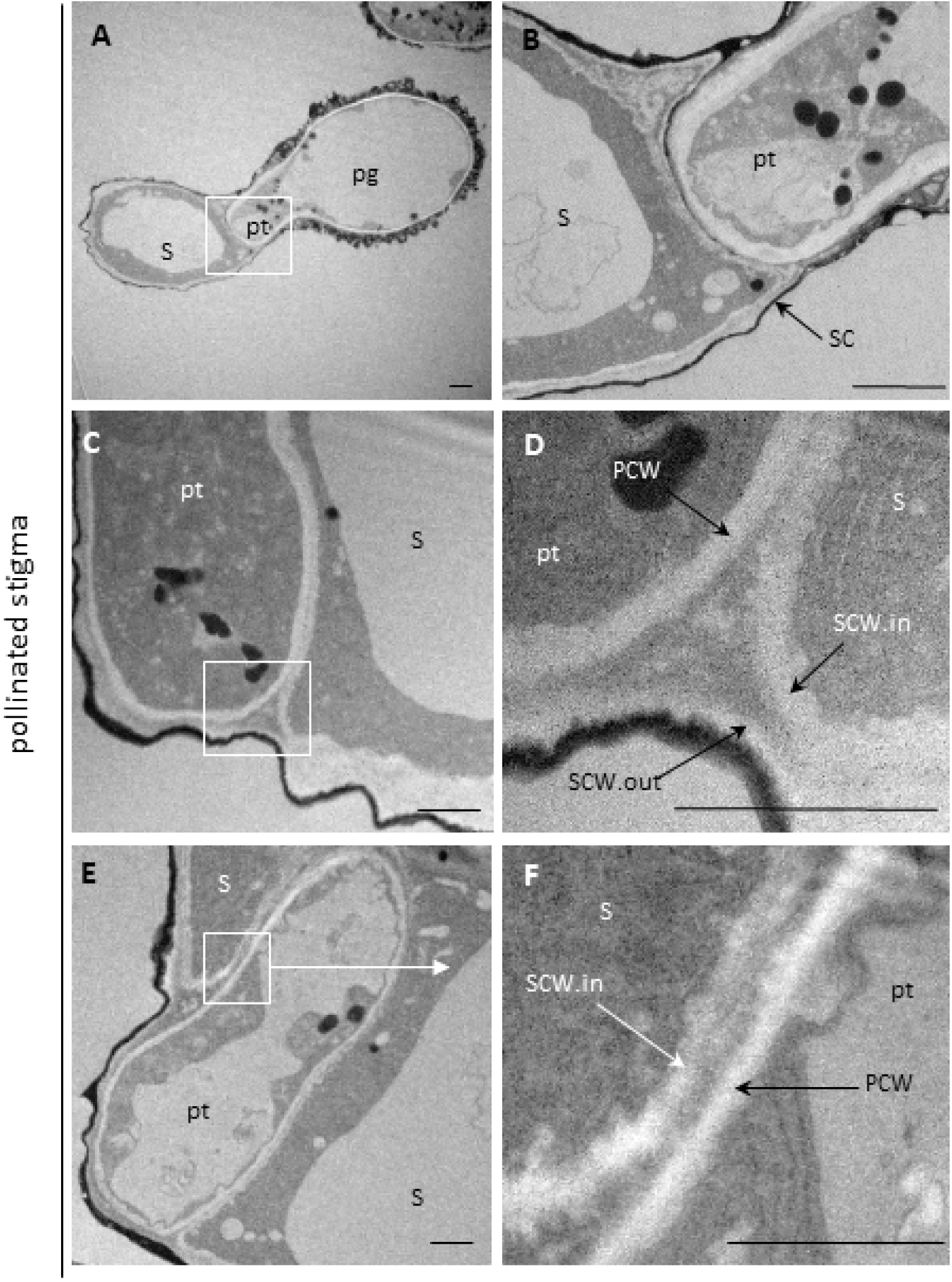
The pollen tube grows within the papilla CW. The images show the pollinated stigma as observed by TEM, 60 map, with general and detailed views in the left and right columns, respectively. A, Transversal section showing the pollen tube (pt) emerging from a pollen grain (pg) and penetrating a stigmatic cell (s). B, close-up view depicted by the white square in A shows stigmatic cuticle (SC) digestion underneath the pollen tube. C, The progressing pollen tube grows between two CW layers of the stigmatic cell (s), the inner layer (SCW.in) and the outer layer (SCW.out), as indicated in the close-up view (D) depicted by the white square in C. Cell walls appear as electron transparent white/light grey layers. E, Transversal section of the pollen tube within the stigmatic CW. F, close-up view depicted by the white square in E, shows the inner stigmatic l surrounds the pollen tube. PCW, pollen CW. Bars in A and B, represent 2µm and, in C to F, 1 µm. Each experiment was repeated at least three times.

Taken together, we found that oomycete hyphae and pollen tubes both penetrate the papilla cuticle. Subsequent growth characteristics depend on different mechanisms mobilized regarding digestion of the stigmatic CW. While the pollen tube is engulfed within the CW, oomycete hyphae digest the two CW layers to grow in contact with the papilla PM.

### Penetration causes different constraints to the papilla surface

To invade a plant tissue, an advancing cell has to exert a pressure on its host (Sanati Nezhad and Geitmann, 2013). To evaluate the mechanical force applied by the invader as it progresses along the stigmatic cells, we quantified papilla deformations induced by hyphae or pollen tubes early after penetration. According to Riglet et al., (2020) two measures were considered: (i) the deformation towards the interior of the papilla (intD), estimated by invagination of the stigmatic PM labeled with the membrane marker LTI6B-GFP and (ii) the deformation towards the exterior of the papilla (extD) estimated on bright field images (Figure 6, A and B). We found that the *P. parasitica* hypha creates a large external deformation during penetration (2.8 µm, Figure 6C, Supplemental Table S1) and slightly deforms the papilla interior (0.6 µm, Figure 6C, Supplemental Table S1). By contrast, pollen tube growth resulted in almost the same extD and intD (2.1 µm and 1.8 µm respectively; Figure 6C; Supplemental Table S1). This quantitative difference could be due to invader diameters, since the pollen tube is significantly larger than the hypha (*i.e.* 4.8 µm and 3.8 µm respectively; Supplemental Figure S1 A and B; Supplemental Table S1). Although the pollen tube and the hypha pierce and penetrate the stigmatic surface, they apparently do not distort the papilla cell to the same extent.

**Figure 6.**
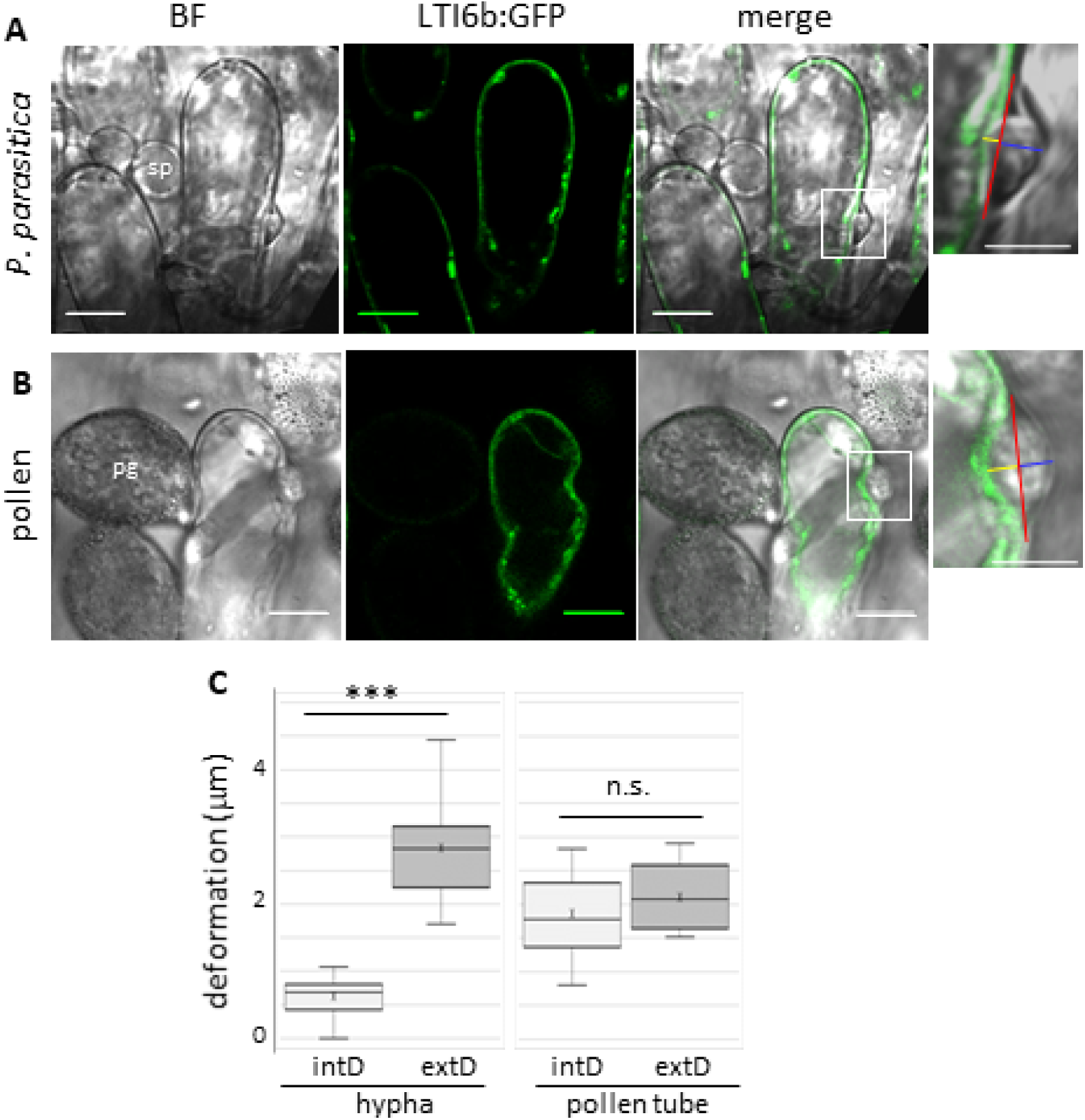
Pathogen and pollen tubes apply different mechanical stresses onto the papilla. Arabidopsis stigma expressing a GFP-tagged PM marker (Lti6b) were infected with *P. parasitica* (A) or pollinated (B), and observed by CLSM at one hai or 30 map, respectively. To quantify the papilla deformation, a red line was drawn on merged images (inset) between the two external points of the deformation. Distances from the red line towards the cuticle (blue line, external deformation, extD) and towards the cytoplasm (yellow line, internal deformation, intD) were determined. Bars represent 10 µm on the full images and 5 µm on the insets. Sp, spore; pg, pollen grain. C, Quantitative analysis of external (extD) and internal (intD) papilla deformations upon infection or pollination, for 21 hyphae or 20 pollen tubes. In the plots, the cross corresponds to the mean value. t-test; *** pVal<0,0005; n.s., not significant.

### Penetration attempts provoke subcellular rearrangements

Vesicle trafficking within infected host cells and remodeling of the cytoskeleton is of crucial importance for plant defense responses (Ruano and Scheuring, 2020; Lu et al., 2023). Similarly, early cellular events associated with pollen acceptance are polarized secretion (Samuel et al., 2009; Safavian and Goring, 2013) and actin reorganization (Iwano et al., 2007; Rozier et al., 2020) oriented towards the pollen grain. We analyzed the trafficking routes and cytoskeletal dynamics triggered in stigmatic cells upon hyphal or pollen tube intrusion, and focused on (i) the trans-Golgi network (TGN), a compartment at the crossroad of secretion and endocytosis (Aniento et al., 2022), (ii) the late endosome (LE), also named multivesicular bodies (MVB), a compartment that passes cargo to the lytic vacuole but can also function in polarized secretion by releasing its internal vesicles in the extracellular space (Aniento et al., 2022), and (iii) the actin network that provides tracks to drive vesicular transport (Geitmann and Nebenführ, 2015). We followed the fate of these cellular components with specific marker lines expressing a GFP-tagged vacuolar ATPase a1 subunit (VHAa1-GFP), a GFP-tagged tandem FYVE domain (GFP-FYVE) and a fusion between the Lifeact peptide and the Venus fluorochrome (Lifeact-Venus), respectively. To observe early penetration stages, pistils were inoculated with *P. parasitica* for one hour or pollinated for 30 minutes and cellular component dynamics was monitored by confocal microscopy. We quantified the intensity of the fluorescence signal in the papilla around the invader penetration site using a homemade Fiji macro. We calculated a difference of fluorescence intensity between the invader penetration site (contact zone) and the surrounding area where there was no penetration (surrounding area). A positive difference [contact-surrounding] suggests a focalisation of the component of interest towards the penetration site. In *P. parasitica-*infected papillae, we detected a significant increase of the TGN-VHAa1 and LE-FYVE fluorescence at the contact zone with the growing hyphae (Figure 7,A and B; Supplemental Figures S2 and S3; Supplemental Table S2). LE-FYVE fluorescence enhanced upon pollen tube intrusion around the penetration site (Figure 7; Supplemental Figure S4, Supplemental Table S2), while the intensity of TGN-VHAa1 labeling at the contact zone was not significantly different from the zone without contact (Figure 7; Supplemental Figure S5; Supplemental Table S2). As control, we did not detect any significant fluorescence variation in non-infected and non-pollinated papillae (Supplemental Figure S6; Supplemental Table S2). In stigmatic cells, the actin cytoskeleton formed a network of fine cables homogeneously distributed along the papillae (Supplemental Figure S6A; Rozier et al., 2020). Upon infection, Lifeact-Venus fluorescence significantly increased at the contact zone, forming a dense and brightly fluorescent patch beneath the growing hypha (Figure 7; Supplemental Figure S7; Supplemental Table S2). Similarly, a significant focal accumulation of actin was detected in pollinated papillae at the contact zone with the growing pollen tube (Figure 7; Supplemental Figure S8; Supplemental Table S2).

**Figure 7.**
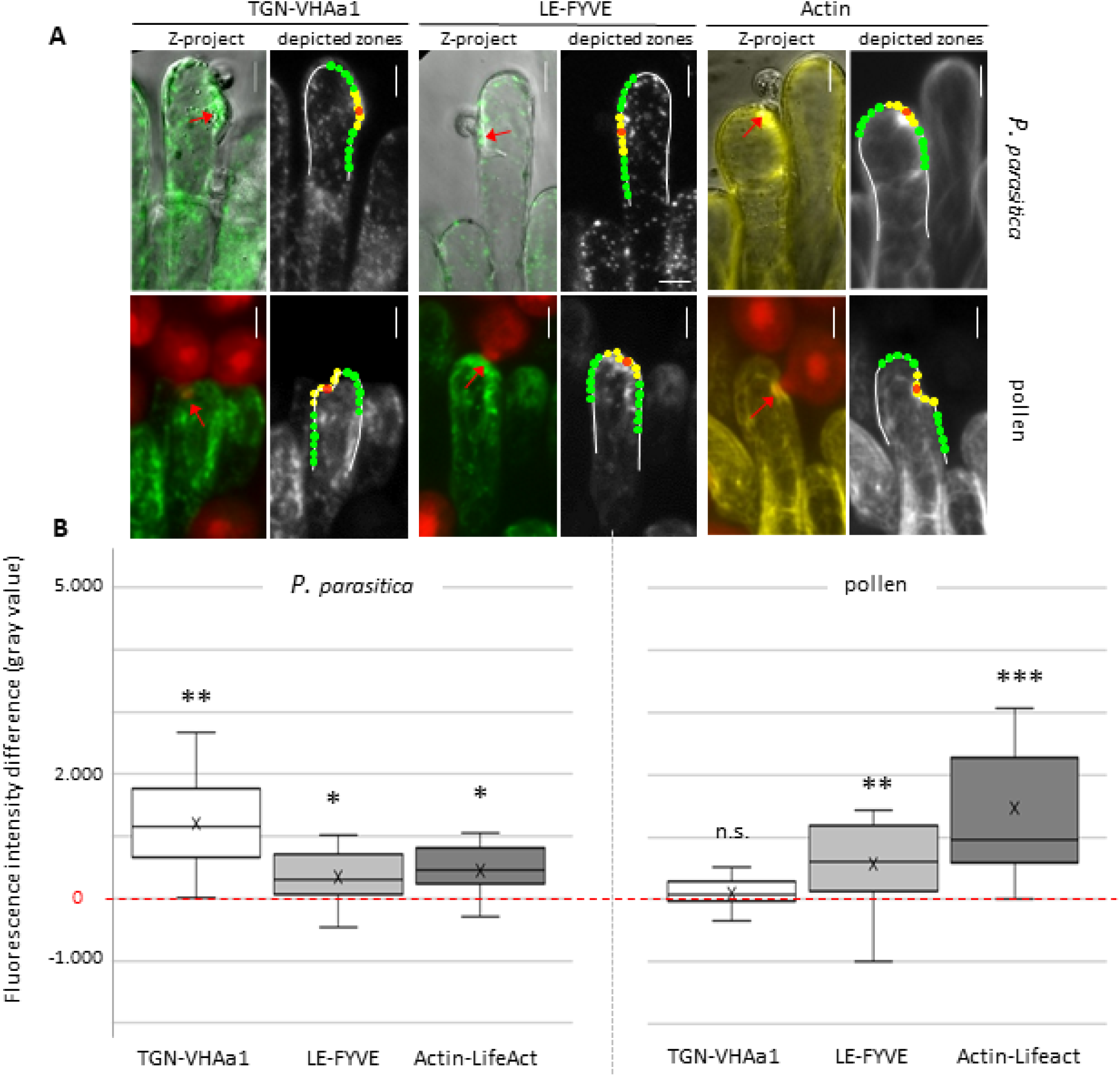
The pathogen and pollen tube trigger both similar and different subcellular rearrangements upon penetration of stigmatic cells. A, Stigmatic cells from fluorescent marker lines for the trans Golgi network (TGN; TGN-VHAa1), the late endosome (LE/MVB; LE-Fyve), and the actin network (Actin-Lifeact) were infected with *P. parasitica* or pollinated. CLSM at one hai or 30 map allowed to visualize the papillae (green or yellow fluorescence) and the invader (bright field for *P. parasitica,* red fluorescence for pollen). The papilla periphery was manually outlined on the obtained images (white lines), and the Fiji macro automatically depicted two zones, (i) a contact zone (yellow circles) including the invader entry point (red arrow/red circle), and (ii) a surrounding zone (green circles). Fluorescence intensities (gray values) were automatically measured in each circle. Bars represent 10 µm. B, Quantification of fluorescence intensity differences between zones (i) and (ii). For each interaction, 15 stigmatic cells on at least four independent stigma were analyzed. In the plots, the cross corresponds to the mean value. Statistical analysis of fluorescence intensity was based on a paired T-test. * pVal<0,05; ** pVal<0,005; *** pVal<0,000.5; n.s., not significant. Detailed measurements are shown on supplemental Figures S2 to S5, S7 and S8.

Dynamic changes in the endomembrane system and the cytoskeleton are still poorly documented upon invasion of the natural target for *P. parasitica* infection; we then extended our comparison to the root epidermis. Similar to infected stigmatic cells, we detected a significant increase of the TGN-VHAa1 and LE-FYVE fluorescence at the contact zone with the growing hyphae in infected root cells (Figure 8A and B; Supplemental Figures S9 and S10; Supplemental Table S2). Such fluorescence focalization was not observed in control root cells (Supplemental Figure S11A and B). Our Fiji macro was not suitable for actin-Lifeact quantification since actin filaments were highly concentrated at the cortical region of the entire root (asterisk in Figure 8C), possibly masking a focal accumulation at the contact zone. We then counted the number of images displaying a large actin-Lifeact focalisation patch at the penetrating hypha tip (red arrow in Figure 8C). The oomycete triggered actin focalization at the contact zone in 11 infected root cells out of 15 (Supplemental Figure S12), while no such fluorescent patches were detected in control cells (Supplemental Figure S11C).

**Figure 8.**
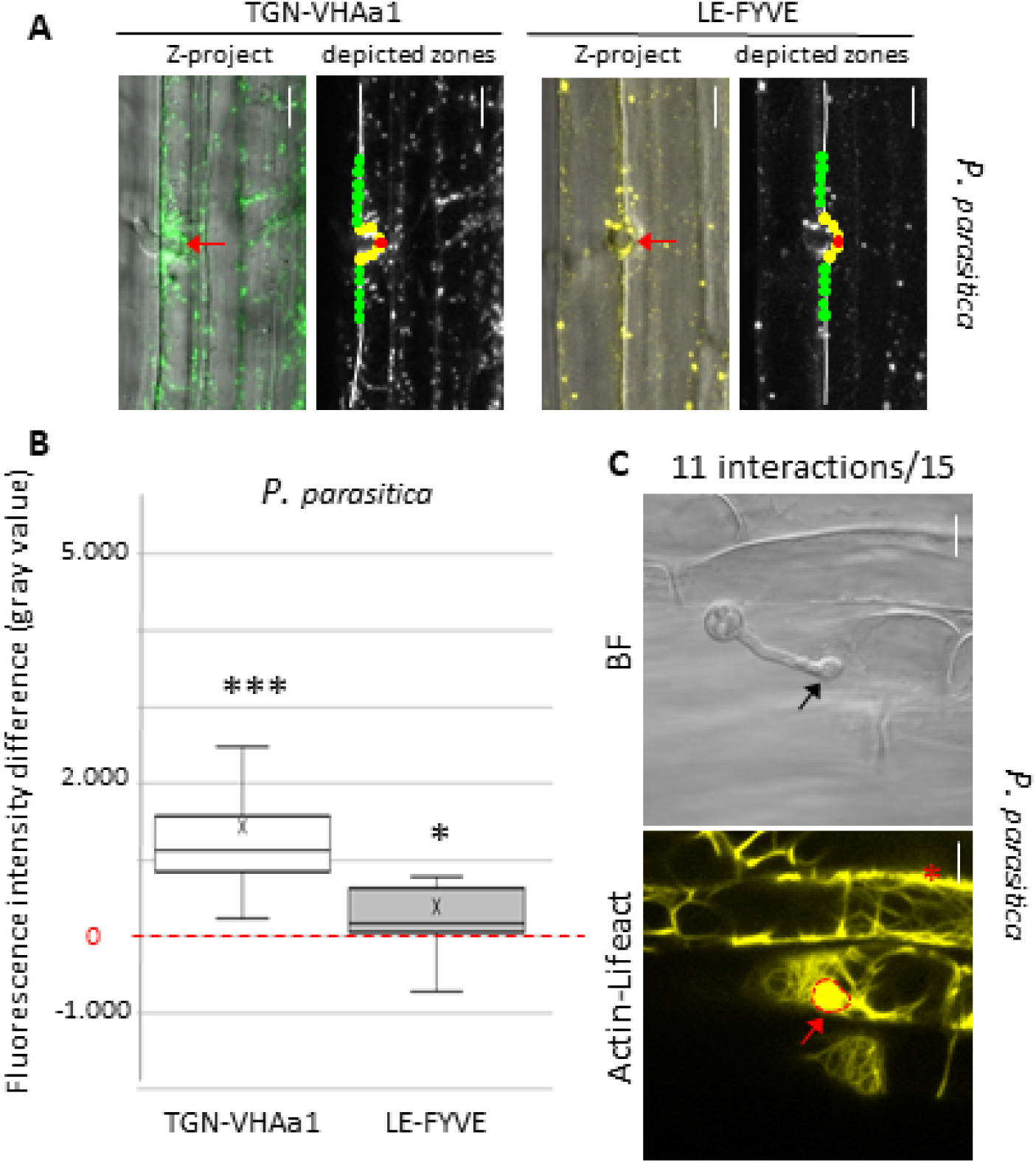
*P. parasitica* triggers reorganization of subcellular components in root cells. Roots from fluorescent marker lines for the trans Golgi network (TGN; VHAa1), the late endosome (LE/MVB; 2xFyve), and the actin network (Actin; LifeAct) were infected with *P. parasitica* and analyzed by CLSM at one hai. A, For the TGN and LE marker lines, fluorescence was quantified using the Fiji macro as described in Figure 7 to determine fluorescence differences between contact zones and surrounding zones B, Quantification of fluorescence differences. For each interaction, 15 root cells on at least 10 independent roots were analyzed. In the plots, the cross corresponds to the mean value. Statistical analysis of fluorescence intensity was based on a paired T-test; * pVal<0,05; *** pVal<0,0005. Detailed measurements are shown on supplemental Figures S9 and S10. C, The Fiji macro was not applicable to quantify actin fluorescence, because the high concentration of actin filaments at the cortical region (asterisk) distorted the quantification of fluorescence. We thus visually determined actin focalization when a fluorescence patch (red arrow, delimited by a red dashed line) was clearly visible at the contact site with the penetrating hyphae (black arrow). Among 15 root cells on 12 independent roots we determined a frequency of 11/15 events of actin focalization at penetration sites (see Supplemental Figure S12). BF, bright field. Bars represent 10 µm.

## Discussion

We aimed at deciphering the response of a single epidermal cell, the papilla, to the intrusion of a pollen tube or the hypha of pathogenic oomycetes from two genera, *P. parasitica* and *H. arabidopsidis*.

Following *P. parasitica* inoculation, we found that swimming zoospores accumulated massively at the stigma and at the elongation zone of the root, suggesting both zones have common features that specifically allow zoospore aggregation. The attraction by root exudates of zoospores from several *Phytophtora* species is not fully elucidated but a wide variety of components, such as carbohydrates, amino-acids and hormones, have already been identified as attractant cues (Bassani et al., 2020; Kasteel et al., 2023). *A. thaliana* belongs to a plant family characterized by a dry stigma without surface exudates (Hiscock and Allen, 2008) suggesting that secreted chemical cues are likely not responsive for spore aggregation at the stigma surface. Electrostatic forces have been suggested to function in both interaction systems. Zoospore accumulation correlates with the natural electrostatic field generated by the root and may explain the asymmetric recruitment of spores (Van West et al., 2002). Mathematical models predicted that the electric field increased near the extremity of the pistil as the flower opened and this may participate in the electrodeposition of pollen grains onto the stigma (Clarke et al., 2017). Although it remains largely unknown how these electrical signals are perceived by the zoospores or the pollen grains, we can speculate that electrostatic forces may be involved in the preferential accumulation of *P.parasitica* spores at the stigmatic surface.

Once at the epidermis surface, filamentous pathogens form appressoria to penetrate the tissue. Early after pollen landing, the foot is formed to strengthen pollen adhesion and to loosen the stigmatic CW to prepare the pollen tube entry (Chapman and Goring, 2010). Thus, both pathogen hypha and pollen tube require specialized structures to penetrate the epidermis. Thereby, the appressorium-like structure differentiated by *P. parasitica* at the outer surface of the stigmatic epidermis (Figures 3 and 4) is likely used to rupture the cuticle and the CW allowing entry within the papillae. Filamentous pathogen appressoria can not only be formed at the host surface but also on many artificial materials such as glass slides and polycarbonate membranes (Bircher and Hohl, 1997; Gaulin et al., 2002). These *in vitro* experiments led some authors to suggest that host factors are not essential for appressorium development. On the other hand, physical and chemical properties of the epidermis, such as surface hardness, hydrophobicity, wax composition and cutin, are strong triggers for the formation of appressoria of fungi and oomycetes (Bircher and Hohl, 1997; Ryder et al., 2022). A hydrophobic cuticle covers the outer surface of the epidermal cell walls of stigma and leaves, but not of roots (Heizmann et al., 2000; Schreiber, 2010). Despite this, we never observed appressorium formation by the leaf pathogen *H. arabidopsidis* on stigma (Figure 2). This may indicate that the papilla either lacks the triggering signals or inhibits the differentiation of *H. arabidopsidis* appressorium. Alternatively, host range specificities may also determine the capacities to infect different plant organs. *H. arabidopsidis* is an obligate pathogen, which exclusively infects *A. thaliana* leaves. By contrast, *P. parasitica* infects more than 72 plant species and produces appressoria on root and leaf tissues under natural conditions (Meng et al., 2014). It is therefore likely that *H. arabidopsidis* depends on specific host stimuli to differentiate appressoria, whereas *P. parasitica* is less selective and develops appressoria on various supports, including stigma.

To overcome the barrier of the plant CW, filamentous pathogens and pollen grains secrete CW-degrading enzymes capable of digesting its main polymers (Kubicek et al., 2014; Robinson et al., 2021). Our TEM analysis of the penetration process (Figures 4 and 5) reveals a major difference between both invaders regarding their ability to digest the stigmatic CW. Whereas the hypha passes through the bilayer papilla wall to grow in between the inner face of the CW and the PM, pollen tube penetration is restricted to the outer wall layer and the tube growth is confined inside the CW. This suggests that the two papilla wall layers may have different chemical properties and their digestion would require diverse cocktails of secreted enzymes; *P. parasitica* would secrete enzymes that can digest both CW layers, whereas pollen have not. Interestingly, heterogeneity of the stigmatic wall has already been suggested (Riglet et al., 2020). In this work, the authors proposed that inner and outer layers have different mechanical properties related to the orientation of cellulose microfibril, which has a strong impact on the pollen tube behavior.

Advancing hyphae and pollen tubes and may exert a mechanical force during invasion to overcome the physical barrier of stigmatic cells. We estimated this compressive force by measuring the papilla deformation (Figure 6). Surprisingly, we found that the pollen tube, while growing engulfed within the rigid CW, deforms the interior of the cell, whereas the hypha, which passes through the CW layers and becomes separated from the cytoplasm only by the stigmatic PM, does it poorly. We may assume that this difference is related to the magnitude of the forces exerted by the two growing structures and/or the different resistance forces generated by the papilla, which may depend on the location or the nature of the invader (Sanati Nezhad and Geitmann, 2013).,Hardham and co-workers showed that touching the surface of the *A. thalian*a cotyledon epidermis with a microneedle produced a rapid actin focalization at the contact point suggesting that actin reorganization is triggered by detection of the mechanical pressure exerted by the invader (Hardham et al., 2008). In stigmatic cells challenged by a hypha or a pollen tube, we detected a similar reorganization of actin around the penetration site (Figure 7). Although the pollen tube deforms the interior of the cell, likely applying a stronger pressure compared to the hypha, a threshold value for the physical forces required to stimulate actin reorganization may have been reached in both interaction systems. Such a mechanical threshold has been proposed to elicit subcellular reorganization in epidermal cells infected with *H. arabidopsidis* (Branco et al., 2017).

Within the cytoplasm of infected roots or stigmatic cells, the growing hypha is surrounded by a membrane envelope labeled with fluorescent markers localized to the PM in non-infected cells (Figure 3). Such membrane envelope were previously described in *A. thaliana* and rice leaves to surround hyphae of the hemibiotrophic fungi *Magnaporthe oryzae* or *Colletotrichum higginsianum* upon infection (Yi and Valent, 2013; Qin et al., 2020). This membrane, called the extra-invasive hyphal membrane (EIHM), is considered as a typical hallmark for the biotrophic phase, required to escape the plant recognition system and to uptake nutrients from the host (Oliveira-Garcia and Valent, 2015; Jones et al., 2021). The EIHM forms a continuum with the plant PM but its composition differs as its content is modified during infection (Qin et al., 2020). To our knowledge such specialized membrane have never been reported for pathosystems involving flower pathogens (Brewer and Hammond-Kosack, 2015; Andargie and Li, 2016; Mondragón-Palomino et al., 2017). The observation of an EIHM-like membrane in papillae cells infected by *P. parasitica*, suggested that the oomycete developed as a biotroph during the first hours after infection, as it does in roots. Whether this envelope is a functional interface and whether its composition differs from the rest of the PM needs further investigation.

The mobilization of TGN and LE/MVB vesicles contributes to the defense mechanisms by delivering defense proteins and antimicrobial compounds to the extracellular space, reinforcing the CW to prevent pathogen entry and participating in the expansion of the host-derived membrane during infection (Bozkurt et al., 2015). Thereby, it was not surprising to find VHA-TGN and FYVE-LE concentrated at the vicinity of *P. parasitica* penetration site in root epidermal cells (Figure 8). Interestingly, TGN and LE focalisation were also triggered by *P. parasitica* in infected stigma (figure 7). Thus, the pathogen is capable of manipulating a non-natural target, the papilla, to hijack the host machinery similarly to what it does in roots. Upon pollination, an intense vesicular trafficking in stigmatic cells is essential to sustain pollen germination and pollen tube growth. Cargos transported by the vesicles are poorly known; they may deliver hydration factors, wall-loosening enzymes and other components to facilitate germination of the incoming grain and penetration of the emerging pollen tube (Doucet et al., 2016). Ultrastructural studies on pollen–stigma interactions in *Brassica napus* identified LE/MVB fused to the stigmatic PM and release of internal vesicles in the extracellular space adjacent to the pollen grain (Safavian and Goring, 2013; Indriolo et al., 2014). In pollinated Arabidopsis stigma, instead of LE/MVB, secretory vesicles, likely originated from the TGN, were detected attached to the stigmatic PM beneath the pollen grain (Safavian and Goring, 2013; Indriolo et al., 2014). In our experiments performed on *A. thaliana*, we did not detect polarized movement of the TGN-VHAa1 compartments during pollen tube penetration. This discrepancy may be related to the methods used between the two studies, confocal imaging and fluorescent probes in our case, chemically fixed material and TEM in Safavian and Goring (2013) and Indriolo *et al*. (2014). Beside, the TGN is a complex compartment at the interface of the secretory and endocytic pathways, divided into subdomains or sub-populations (Aniento et al., 2022) and deciphering its implication in pollination would require additional live-cell imaging with a large collection of fluorescent-tagged markers for endomembrane compartments. Nethertheless, our study highlighted that LE/MVB trafficking could be implicated in pollen acceptance in *A.thaliana*, in addition to the conventional secretory pathway suggested by Goring and colleagues (Safavian and Goring, 2013; Indriolo et al., 2014).

In conclusion, we identified an epidermal cell that enables the in-depth comparative analysis of how the same plant cell responds to invading hyphae of a pathogen and to the beneficial process of pollination. Certain of the subcellular changes triggered in stigmatic cells upon *P. parasitica* infection are also activated in responses to pollen (LE trafficking and actin remodeling). Our work supports the long-standing assumption that common or evolutionary related host-encoded functions exist between plant defense and pollen recognition. Besides, we highlighted cellular events specific for each type of interaction (TGN mobilization, formation of an EIHM-like membrane) that may be required for an appropriate response likely depending on the chemical nature of the invader and/or the established entry strategy (*i.e*. appressorium formation and complete CW digestion *vs* foot formation and partial CW digestion). The engulfing of the pollen tube within the stigmatic CW remains quite enigmatic. Little is known about the composition of the papilla wall, except that the bilayered structure is not found in other epidermal cells. From our data, it is tempting to speculate that limitation of the CW digestion to the outer layer and constrained growth of the pollen tube inside the wall, may represent specialized adaptations to discriminate a pollen tube from an unwanted invasive agent, such as a pathogen. It is likely that more crosstalks exist between infection and pollination. The interaction system we have developed may provide a framework for the further exploration of how an epidermal cell senses and responds to an invader in order to adjust the most relevant responses.

## Materials and methods

### Biological material and culture conditions

All *Arabidopsis thaliana* lines were in the Col-0 background and grown in growth chambers under long-day conditions (16h light/8h dark at 21°C/19°C with a relative humidity around 60%). Three sets of Arabidopsis marker lines were used; (i) For expression in stigmatic cells, we used the Brassica pSLR1 promoter (Rozier et al., 2020). The pSLR1-LTI6b:GFP and the pSLR1-Lifeact:Venus lines were previously described (Rozier et al., 2020). We used the Gateway® technology (Life Technologies, USA; http://www.thermofisher.com, (Karimi et al., 2002) to generate the pSLR1-GFP:2xFyve construction. (ii) to control expression in root cells, two ubiquitous promoters, p35S and pUbiquite10, were used; these promoters are poorly active in papillae. We generated the p35S-*At*PIP2A:RFP construction in the binary vector, pm-rk (Nelson et al., 2007), using the Gateway® technology. The pUbiquitine10-Citrine:2xFyve and the pUbiquitine10-Lifeact:YFP lines were previously described (Simon et al., 2014; Doumane et al., 2021). The pVHAa1-*At*VHAa1:GFP line was described previously (Dettmer et al., 2006); the *VHAa1* promoter is active in both stigmatic and root cells. (iii) The pACT11-RFP and the pLAT52-GFP lines, expressing a cytoplasmic fluorescent marker in pollen grain and tube, were previously described (Rotman et al., 2003; Rozier et al., 2020).

*P. parasitica* Dastur isolate INRA-310 was maintained in the Phytophthora collection at INRAE, Sophia Antipolis, France. The growth conditions and zoospores production were previously described (Galiana et al., 2005). The *P. parasitica* transformant (pCL380-GFP:GUS) expressing a GFP:GUS fusion protein was previously described (Attard et al., 2014). The *H. arabidopsidis* isolate Noco was transferred weekly onto the susceptible accession Col-0 as described (Hok et al., 2014). Inoculated plants were kept in a growth cabinet at 16°C for 6 days with a 12 h photoperiod.

### Oomycetes pathogen assays and histochemical analysis

Pathogen assays with the *P. parasitica* and *H. arabidopsidis* isolates on roots and leaves, respectively, were performed as previously described (Hok et al., 2014; Le Berre et al., 2017). To infect pistil tissues with *H. arabidopsidis*, Arabidopsis leaves with the sporulation oomycete on their surface were gently rubbing over the pistil surface of manually opened flower buds (late stage 12; Smyth et al., 1990). Alternatively, spores were applied in solution (5×10^5^ zoospores/ml) directly on the stigma surface. Inoculated pistils were observed by confocal microscopy in a period of 4h to 24h after inoculation. Manually opened floral buds or naked pistils (late stage 12; Smyth et al., 1990), were dipped in an aqueous suspension of *P. parasitica* zoospores (5×10^5^ zoospores/ml) obtained from the strain pCL380-GFP:GUS (Attard et al., 2014). In a period of 3h to 24h after infection, the GUS reporter activity staining in plant tissues was performed as previously described (Hok et al., 2014).

### Pollination assay and aniline blue staining

Pistils (late stage 12; Smyth et al., 1990) were emasculated and pollinated with mature pollen. Six hours after pollination, stigmas were fixed in acetic acid 10%, ethanol 50% and stained with Aniline Blue for epifluorescence microscopy observation as previously described (Rozier et al., 2020).

### Transmission Electron microscopy

Pistils (late stage 12; Smyth et al., 1990) were emasculated and inoculated with *P. parasitica* for three hours or pollinated with mature pollen for 60 minutes. Roots were inoculated with *P. parasitica* for three hours. Pollinated or inoculated tissues were fixed in a solution containing 2.5% glutaraldehyde and 2.5% paraformaldehyde in 0.1 M phosphate buffer (pH 7.2) and after four rounds of 30 min vacuum, they were incubated in fixative for 12 hours at room temperature. Pistils or roots were then washed in a phosphate buffer and further fixed in 1% osmium tetroxide in 0.1 M phosphate buffer (pH 7.2) for 1.5 hours at room temperature. After rinsing in phosphate buffer and distilled water, samples were dehydrated through an ethanol series, impregnated in increasing concentrations of SPURR resin over a period of three days before being polymerized at 70°C for 18 h, sectioned (65 nm sections) and imaged at 80 kV using an FEI TEM tecnaiSpirit with 4 k x 4 k eagle CCD.

### Confocal Laser Scanning Microscopy (CLSM)

Pistils (late stage 12; Smyth et al., 1990) were emasculated and inoculated with *P. parasitica* for one hour or pollinated with mature pollen for 30 minutes. Roots were inoculated with *P. parasitica* for one hour. Pollinated pistils were observed with a Zeiss microscope (Zeiss 800 or AxioObserver Z1 equipped with a spinning disk module) with a 40x objective. Oomycetes infected tissues (stigma or root) were observed with a Zeiss 880 confocal microscope with a 63x objective. Venus, Citrine, YFP and GFP were excited at 488 nm and fluorescence detected between 500 and 550 nm. RFP was excited at 561 nm and fluorescence detected between 550 and 600 nm. Stigmas or roots were imaged every 0.4 µm, encompassing the entire volume of the stigma or half of the root thickness, using z-stack confocal protocol. Pictures were taken with detector settings optimized for no pixel saturation.

### Fluorescence quantification

All image processing, image analysis and fluorescence measurements were done using the ImageJ/Fiji program (Schindelin et al., 2012).

To quantify fluorescence intensity at the contact site with the invader, we used a homemade Fiji macro. From the serial confocal images, we generated an average intensity projection (Z project). We manually choose one slide from the stack which corresponds to the focus plan of the contact site with the invader. On this selected slide, we manually drew the stigmatic cell periphery and designated the invader entry point. Then, we indicated the contact area length (ROI zone Length). From numerous image observations, we defined this contact area as 16 µm (Supplemental Figure S1 C-E). Next, the macro automatically depicted two zones, the contact and the surrounding zones, with a series of circles of fixed diameter (ROI zone thickness set at 2.6 um). We estimated that 2.6 µm was an appropriate dimension compared to the papilla sizes and the contact area length. The contact zone included five to seven circles depending on its shape (straight or curved. The surrounding zone contained twice as many circles as the contact zone equally distributed from each side of the contact area. Fluorescence intensity was measured in each circle by the Fiji script, given as gray values and reported in an Excel file. The mean fluorescence in contact and surrounding zones was calculated, then, a fluorescence difference [contact-surrounding] was applied. For control stigmatic cells (non infected or non pollinated), as there was no invader entry point, we introduced zero for the ROI zone Length, 2.6 um for ROI zone thickness and defined an arbitrary contact zone of six circles always positioned at the same distance from one extremity of the drawn papilla periphery. This Fiji macro is available on demand.

For fluorescence quantification in root cells and vesicular marker lines, we followed the same procedure except that we manually outlined one root edge. For actin fluorescence, quantification using the Fiji macro was not possible. We then counted the number of images displaying a large actin actin focalization when a fluorescence patch was clearly visible at the contact site with the penetrating hyphae.

Statistical analyses of fluorescence intensity at the contact site with the invaders were based on the paired sample *t*-test. The statistical analysis was carried out on control and inoculated or pollinated cells (n =15).

### Deformation and diameter measurements

Pistils (late stage 12; Smyth et al., 1990) expressing a GFP-tagged PM marker (LTI6b) were emasculated and inoculated with *P. parasitica* or pollinated with mature pollen. Stigma were observed under CLSM one hai or 30 map respectively. Internal (IntD) and external (ExtD) papilla deformation were measured at the penetration site with the invaders as described (Riglet et al., 2020). The statistical analysis compared IntD and ExtD (n= 21 hyphae, 20 pollen tubes) and were based on umpaired *t*-test. On the same LTI6bGFP images, we measured two perpendicular diameters of the hypha or pollen tube and calculated a mean diameter. The statistical analysis compared both diameters (n= 21 hyphae, 20 pollen tubes) and were based on umpaired *t*-test.

## Supplemental data

**Supplemental Figure S1.** Invader features.

**Supplemental Figure S2.** Pattern of TGN-VHAa1 compartments in stigmatic cells in response to *P. parasitica*.

**Supplemental Figure S3.** Pattern of LE-FYVE compartments in stigmatic cells in response to *P. parasitica*.

**Supplemental Figure S4.** Pattern of LE-FYVE compartments in stigmatic cells in response to pollen tubes.

**Supplemental Figure S5.** Pattern of TGN-VHAa1 compartments in stigmatic cells in response to pollen tubes.

**Supplemental Figure S6.** Subcellular arrangement of the TGN, the LE, and the actin network in control stigmatic cells as observed by CLSM.

**Supplemental Figure S7.** Pattern of the actin cytoskeleton (Actin-Lifeact) in stigmatic cells upon infection with *P. parasitica*.

**Supplemental Figure S8.** Pattern of the actin cytoskeleton (Actin-Lifeact) in stigmatic cells in response to pollen tubes.

**Supplemental Figure S9.** Pattern of TGN-VHAa1 compartments in root cells in response to *P. parasitica*.

**Supplemental Figure S10.** Pattern of LE-FYVE compartments in root cells in response to *P. parasitica*.

**Supplemental Figure S11.** Subcellular arrangement of the TGN, the LE, and the actin network in control root cells, as observed by CLSM.

**Supplemental Figure S12 :** Dynamics of the actin cytoskeleton (Actin-Lifeact) in root cells in response to infection with *P. parasitica*, as analyzed by CLSM at one hai.

## Acknowledgments

We thank all the members of the Cell Signaling and Endocytosis (SiCE) group (Laboratoire de Reproduction et Développement des Plantes, ENS Lyon, France) and the Interactions Plantes-Oomycètes (IPO) team (Institut Sophia Agrobiotech, Sophia Antipolis, France) for fruitful discussions. We thank Y. Jaillais for his comments on the manuscript. We thank P. Bolland, A Lacroix, J. Berger (RDP) for plant care. We thank for the microscopy facility, the “Institut Sophia AgroBiotech” (PlantBIOs) and the “Institut de Pharmacologie Moléculaire et Cellulaire”, both part of the “Microscopie Imagerie Cytométrie Côte d’Azur” GIS IBiSA labeled platform, and the PLATIM of the SFR Biosciences Gerland-Lyon Sud. We thank C. Lionnet at the PLATIM and F. Brau at the “Institut de Pharmacologie Moléculaire et Cellulaire”. We thank the Bordeaux imaging center for TEM imaging and analysis.

## Author contribution

LR was responsible for all experiments and analysis performed for pollination. SH developed the pathosystem for papilla infection and performed the confocal imaging. LBV performed the image acquisition by SEM. NM performed and analyzed *P. parasitica* root and pistil infections. JL performed and analyzed *P. parasitica* root infections. VA and HK performed *H. arabidopsidis* leaf and pistil infections. VB designed the Fiji macro. LR, SH, TG, HK, MG, IFL and AA designed the study; LR, IFL and AA wrote the manuscript. All the authors contributed to the discussion, reviewed and edited the manuscript. The authors read and approved the final manuscript.

